# Unexpected long-term retention of subcutaneous beeswax implants and additional notes on dose and composition from four testosterone implant studies

**DOI:** 10.1101/2022.06.30.498255

**Authors:** Jordan Boersma, Alexandra McQueen, Anne Peters, Joseph F. Welklin, Sarah Khalil, René Quispe, Wolfgang Goymann, Hubert Schwabl

## Abstract

Experimental manipulations of testosterone have advanced our understanding of the hormonal control of traits across vertebrates. Implants are commonly used to supplement testosterone and other hormones to organisms, as they can be readily scaled to produce desired hormone levels in circulation. Concerns about pharmacological (i.e. unnatural) doses of traditional silastic implants led to innovation in implant methods, with time-release pellets and beeswax implants proposed as solutions. A study comparing silastic, time-release pellets, and beeswax implants found the latter to be most effective in delivering a physiologically relevant dose. One proposed advantage to subcutaneous beeswax implants is that they are expected to degrade within the body, thus removing the obligation to recapture implanted individuals in the field. However, few studies have reported on dosage and no published literature has examined the assumption that beeswax implants readily degrade as expected. Here we present time-release androgen data in relation to implants containing varying levels of testosterone from four separate implant studies. In addition, we report long-term persistence of subcutaneous implants, including two cases of implants being retained for > 2 years. Finally, we offer recommendations on the composition and implementation of beeswax implants to aid the pursuit of minimally invasive and physiologically relevant manipulations of circulating hormones.

## 1. Introduction

The androgen testosterone has pleiotropic effects on morphology and behavior, and can integrate multiple inter-linked traits underlying phenotypes (Cox et al., 2016; Fuxjager et al., 2018; Hau, 2007; Hau and Goymann, 2015; Lipshutz et al., 2019). Much of what we know about the functions of testosterone has been learned through manipulations whereby hormone signaling is suppressed or enhanced in captive or wild vertebrates. Illustrating that a trait changes when testosterone signaling is blocked or enhanced indicates it is under androgenic control. Early studies in captive animals used castration of males to remove natural testosterone production to determine if the trait is lost and then supplemented testosterone to determine if the trait is recovered (Adkins-Regan, 2005; Adkins, 1977, 1975; Balthazart et al., 1983; Berthold and Quiring, 1944). These studies typically delivered a pharmacological (i.e. unnatural) dose of testosterone, for example via injection of the hormone, and illustrated which traits are sensitive to the hormone. However, to determine whether a trait is regulated by testosterone under natural conditions, one must deliver a physiologically relevant dose (Fusani, 2008; Goymann and Dávila, 2017; Quispe et al., 2015).

For several decades subcutaneous testosterone implants using silastic tubing have been used in wild animals to determine which traits are sensitive to enhanced testosterone circulation (Balthazart et al., 1983; Boersma et al., 2020; Cordero, 2008; Enbody et al., 2022; Fusani, 2008; Lahaye et al., 2014; Lindsay et al., 2016, 2011; Muck and Goymann, 2018; Sandell, 2007). Yet, two challenges of testosterone implants endure today: 1) ensuring the dose is physiologically relevant, and 2) designing minimally invasive implants that do not require recapture and removal. Scaling testosterone implants to deliver a physiological dose in wild organisms is challenging because one must first establish natural variation in testosterone circulation across and within individuals and then finely scale implants to mimic natural circulation. Often when implants are scaled to deliver a physiological dose of testosterone, sampling shortly after implantation reveals pharmacological peaks (Goymann and Wingfield, 2014; Quispe et al., 2015). In bird studies, testosterone contained within silastic tubing has been the most common form of implant (Balthazart et al., 1983; de Jong et al., 2017; Fusani, 2008; Gerlach and Ketterson, 2013; Lindsay et al., 2011; Moore, 1982; Peters, 2007; Podmokła et al., 2018; Quispe et al., 2015; Siefferman et al., 2013; Sperry et al., 2010). However, these implants require removal, which can be a major challenge for studies of wild birds. Time-release pellets are a more recent innovation that do not require removal because the matrix containing the hormone is thought to be steadily digested (Fusani, 2008; Quispe et al., 2015). An initial study found these commercially available pellets also seem to deliver a more consistent dose over time when compared to silastic implants, thus allowing appropriately scaled implants to maintain a physiological dose absent pharmacological peaks of testosterone (Fusani, 2008). However, a subsequent study found testosterone peaked beyond physiological levels for 1-2 weeks before leveling off for the rest of the 90-day manipulation period (Edler et al., 2011). Another drawback to these pellets is that they are relatively expensive and cannot be readily scaled by experimenters as doses are set by the manufacturer (Fusani, 2008).

Implants of beeswax mixed with peanut oil as carrier matrix for the hormone are an attractive alternative to silastic implants and commercial time-release pellets, as they are cost-effective, can be scaled to deliver desired doses, and are thought to be readily digestible by the implanted organism (Quispe et al., 2015). When comparing such implants to time-release pellets and silastic implants, Quispe et al. (2015) found for testosterone supplementation that they were most effective at delivering a consistent physiological dose for less than 2 weeks. Since this initial beeswax/oil implant study, these implant methods have been employed in several birds (Beck et al., 2016; Boersma et al., 2020; Khalil et al., 2020; McQueen et al., 2021) and at least one mammal species (Matas et al., 2020). However, information on time-release of varying dosage is sparse apart from one corticosterone beeswax implant study by Beck et al. (2016), which reported that 12 of 24 implanted birds had dissolved implants 35 days after implantation. Here we show androgen levels resulting from three testosterone dosage levels of implants composed of a beeswax/peanut oil matrix and administered to four bird species. In addition, we report on unexpected long-term retention of the beeswax and peanut oil matrix and offer considerations for future innovation.

## 2. Materials and Methods

### 2.1 Study timeline and general framework

We combined data collected across separate studies spanning from 2015 – 2019 in four Australian endemic passerine species (Table 1): a) male red-backed fairy-wren (*Malurus melanocephalus*; Khalil et al., 2020), b) male superb fairy-wren (*Malurus cyaneus*; McQueen et al., 2021), c) male and female white-shouldered fairy-wren (*Malurus alboscapulatus*; Boersma et al., 2020), and d) captive female zebra finch (*Taeniopygia guttata*; Goymann and Schwabl, unpublished data). Each study used similar beeswax and peanut oil implants following Quispe et al. (2015). Implants were prepared in batches of 200 by vortexing melted beeswax and peanut oil, loading into a 1 ml plastic syringe, then dispensing a string of the hardened beeswax/oil mixture and cutting into 20mg fragments. Control implants contained only beeswax and peanut oil, and testosterone implants had a dose of crystalline testosterone dissolved in 100% ethanol mixed in. Proportions of beeswax and peanut oil varied slightly as did the dose of dissolved testosterone across studies (Table 1). We used the same batch of implants for red-backed and most white-shouldered fairy-wrens though we slightly modified the fraction of beeswax to peanut oil following an initial white-shouldered fairy-wren pilot study. Implants were stored in phosphate buffered saline (PBS) until deployment. We inserted implants subcutaneously above the left thigh and sealed the incision site with VetBond™ (3M).

**Table 1.** Composition of implant matrix (proportion of beeswax and peanut oil), testosterone dose, and number of implanted individuals across four bird species. Number of implanted birds shows total (number of individuals implanted with either a testosterone or control implant. Different proportions and sample sizes in white-shouldered fairy-wren show information across two study years. Captive zebra finches were given one of two testosterone doses.

| Species | % beeswax | % peanut oil | Testosterone dose | # implanted | Study |
| --- | --- | --- | --- | --- | --- |
| red-backed fairy-wren | 75 | 25 | 0.5 mg | 39 | Khalil et al., 2020 |
| superb fairy-wren | 80 | 20 | 1 mg | 47 | McQueen et al., 2021 |
| white-shouldered fairy-wren | 80 \| 75 | 20 \| 25 | 0.5 mg | 16 \| 22 | Boersma et al., 2020 |
| zebra finch | 80 | 20 | 1 mg; 2 mg | 16 | unpublished |

### 2.2 Androgen sampling and measurement

In most cases a blood sample was taken on the day of implantation (day 0) for later hormone analysis, and post-implantation blood sampling regimen varied across studies (Figures 1 and S1). Male red-backed and female white-shouldered fairy-wrens were consistently sampled 6 – 14 days after implantation to collect pin feathers molting in response to testosterone (Boersma et al., 2020; Khalil et al., 2020). Superb fairy-wrens were sampled opportunistically for several months after implantation, and captive zebra finches were resampled 1, 12, and 19 days after initial implantation. Blood samples were stored on ice in the field for red-backed and superb, but not white-shouldered fairy-wren, and for captive zebra finches blood samples were immediately processed after bleeding. Across studies, blood samples were spun in a centrifuge to separate plasma for androgen analysis. Plasma was frozen in all studies except in white-shouldered fairy-wren, in which plasma was transferred to a 1.5 ml Eppendorf™ containing 400 μl of 100% ethanol following the procedures of Goymann et al. (2007).

**Figure 1.**
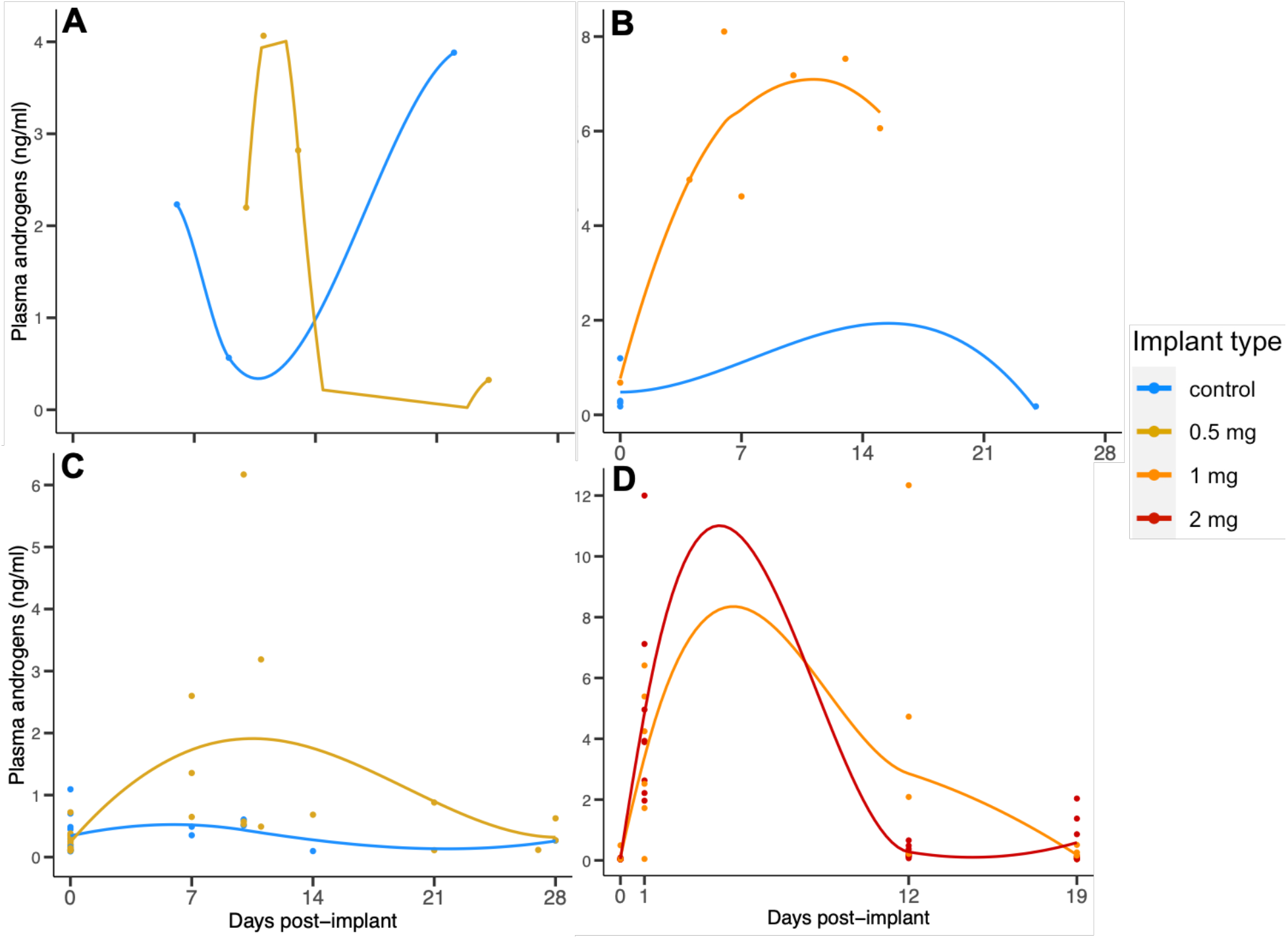
Plasma androgen concentrations (ng/ml) measured within 30 days of implantation with a control (blank) or testosterone (approx. doses = 0.5mg, 1 mg, and 2mg) implant across four species. **A**. red-backed fairy-wren (RBFW; n = 7 males), **B**. superb fairy-wren (SUFW; n = 12 males), **C**. white-shouldered fairy-wren (WSFW; n = 9 males, 43 females), **D**. captive zebra finch (ZEFI; n = 63 females). Individual hormone levels starting on day of implantation (day 0) are shown with loess lines showing the trend for each implant type.

Red-backed and white-shouldered fairy-wren samples were assayed for total androgens following the same radioimmunoassay protocol in a lab at Washington State University (full details in Enbody et al., 2018; Lindsay et al., 2009). Zebra finch plasma was assayed for total androgens using an established radioimmunoassay protocol at the Max Planck Institute for Ornithology, Seewiesen following the procedures detailed in Apfelbeck and Goymann (2011). The detection limit of the zebra finch assay was 0.43pg/tube. All zebra finch samples (N=64) were assayed in one assay with an intra-assay coefficient of variation of 1.9%. In all but superb fairy-wrens, hormone levels reflect total androgens due to using an antibody (Wien Laboratories T-30003, Flanders, NJ, USA) that cross-reacts with closely related steroids, particularly 5α-dihidrotestosterone (DHT). In the superb fairy-wren study, an enzyme immunoassay kit was used to measure testosterone following Crino et al., (2018). Androgen measurements that were below the standard curve in were assigned a value of 1.95 pg/tube in white-shouldered and red-backed fairy-wren (n = 30 samples), and 177.34 pg/ml in superb fairy-wren (n = 5 samples). The 3 non-detectable zebra finch samples were assigned individual detection limits that depended on the respective amount of plasma and extraction recovery (values were 23.3, 27.9, and 40.7 pg/ml). Due to variability in sample storage and assay protocols we do not provide any comparative analyses of androgen levels across species.

### 2.3 Long-term implant retention

State of implants was assessed upon opportunistic recapture of 88 individual red-backed, superb, and white-shouldered fairy-wrens. We checked for signs of infection and noted whether implants were present. No signs of infection were noted in superb or red-backed fairy-wrens, and we noted one case of local inflammation in white-shouldered fairy-wrens. All individuals were monitored for several weeks after implanting and showed no signs of irritation or impairment. In rare cases (n = 3 across species), failure to seal the incision site properly led to implants being lost within several days of initial capture. Post-implant recaptures ranged from 18-400 days in red-backed fairy-wren, 6 – 499 days after implantation in superb fairy-wren, and 7 – 828 days in white-shouldered fairy-wren.

### 2.4 Statistical analysis

We first filtered all androgen samples to only include individuals sampled within 4 weeks of implanting with a testosterone or control implant. For all three fairy-wren species, we assessed whether testosterone-implanted individuals had elevated androgens during this time period using generalized additive models in R (<www.r-project.org>) version 3.6.1, using package mgcv (Wood, 2011). A continuous days post-implant variable was included as a smoothed fixed effect, with testosterone and control-implanted individuals separated. In white-shouldered fairy-wren we included sex as a fixed effect due to having samples from males and females. For zebra finch samples, we compared each time point using package lme4 (Bates et al., 2015) to build a linear mixed model with dose (1 mg or 2mg) and sampling day (0, 1, 12, and 19), and the interaction between dose and sampling day as fixed effects. Individual ID was included as a random effect in white-shouldered fairy-wren and zebra finch due to repeated measures.

### 2.4 Ethical note

For each species testosterone implant and capture protocols were approved by ethical oversight committees. Red-backed fairy-wren work was approved by the Tulane University Institutional Animal Care and Use Committee (IACUC 2019-1715), Cornell University IACUC (2009-0105), Washington State University IACUC (ASAF #04573), the James Cook University Animal Ethics Committee (A2100) and under a Queensland Government Department of Environment and Heritage Protection Scientific Purposes Permit (WISP15212314). Superb fairy-wren work was conducted with approval from the Monash University Animal Ethics Committee (BSCI/2013/10, BSCI/2016/03), Department of Environment, Land, Water and Planning (permit no. 10007370), and the Australian Bird and Bat Banding Scheme (authority nos. 2230, 3288). For white-shouldered fairy-wren, implant work was approved under IAUCUC protocol #0395 and ASAF #04573, and the Conservation and Environment Protection Authority (CEPA) in Papua New Guinea. Finally, housing and implants in captive zebra finch was conducted under the auspices of the Government of Upper Bavaria (permit no. 55.2-1-54-2531-107-10).

## 3. Results

Testosterone implantation led to marginally non-significant elevation of androgens in male red-backed (F_2.51_ = 14.86, *P* = 0.08) and testosterone in male superb fairy-wren (F_2.28_ = 3.83, *P* = 0.06), likely due to low sample sizes within 2 weeks of implantation (Figure 1a,b). Androgens were elevated following implantation in white-shouldered fairy-wren (F_3.35_ = 8.71, *P* < 0.001), with the highest levels measured 7-11 days after implanting (Figure 1c); and sexes did not differ (F_1_ = 0.39, *P* = 0.53). In captive female zebra finch (Figure 1d), 1 mg and 2 mg doses did not differ in plasma androgens (X^2^ = 0.21, DF = 1, *P* = 0.54), but there was a marginally non-significant interaction between dose and sampling day (X^2^ = 6.49, DF = 3, *P* = 0.09). Summary statistics for zebra finch androgens across doses and sampling days are shown in Table 2a. Testosterone was significantly elevated relative to pre-implant (day 0) levels on days 1 and 12, but not 19 (Table 2b). Androgens sampled from 31 days to the end of sampling in all three fairy-wren species are shown in Figure S1. The minimum percentage of implanted individuals who retained implants across the fairy-wren studies is shown in Figure 2.

**Table 2.** Plasma androgen concentration of zebra finch by dose and sampling day **(a)**, and post-hoc Tukey comparisons of sampling day **(b)**. Bold values reflect significant different concentrations in androgens between sampling days.

**a) Plasma androgens by ZEFI implant dose**
| Dose | Day | Mean (ng/ml) | Std. dev. |
| --- | --- | --- | --- |
| 1 mg | 0 | 0.10 | 0.16 |
| 2 mg | 0 | 0.04 | 0.02 |
| 1 mg | 1 | 3.78 | 2.42 |
| 2 mg | 1 | 4.84 | 3.34 |
| 1 mg | 12 | 2.85 | 4.51 |
| 2 mg | 12 | 0.28 | 0.21 |
| 1 mg | 19 | 0.18 | 0.16 |
| 2 mg | 19 | 0.58 | 0.76 |

**b) Post-hoc comp. across ZEFI androgen sampling days**
| Comparison | $\beta$ | SE | z | P |
| --- | --- | --- | --- | --- |
| 0 vs. 1 | 3.68 | 1.10 | 3.36 | **<0.01** |
| 0 vs. 12 | 2.76 | 1.10 | 2.51 | **<0.05** |
| 0 vs. 19 | 0.09 | 1.10 | 0.08 | 0.94 |
| 1 vs. 12 | -0.93 | 1.13 | -0.82 | 0.83 |
| 1 vs. 19 | -3.60 | 1.13 | -3.17 | **<0.01** |
| 12 vs. 19 | -2.67 | 1.13 | -2.36 | 0.06 |

**Figure 2.**
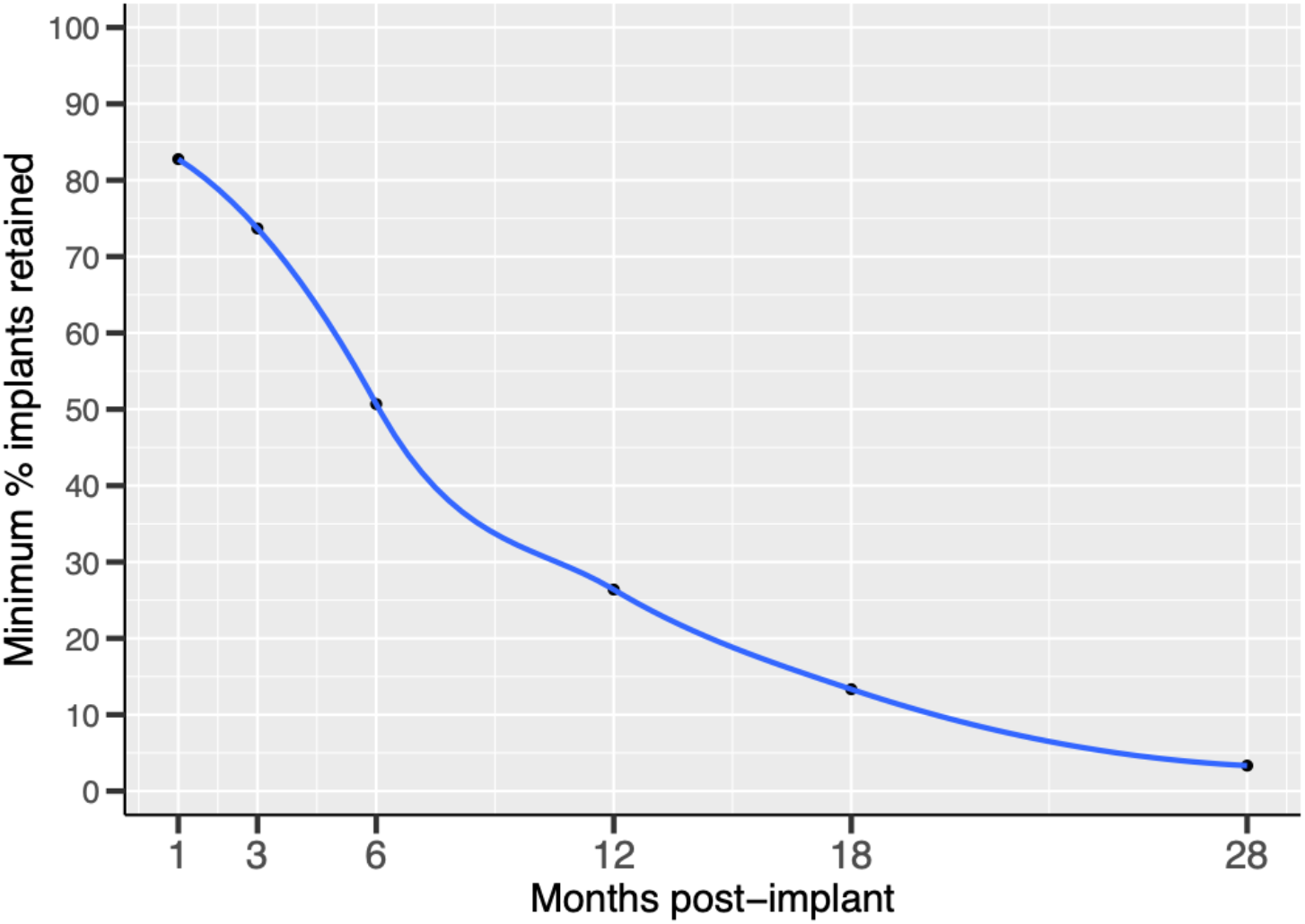
Minimum percentage of implants retained across months of study of free-living fairywrens. Implant retention was assessed on recapture of 88 individual red-backed, superb and white-shouldered fairy-wrens. Only 16 of 88 implanted individuals were confirmed to have fully dissolved implants during the study. Minimum percentage of implants retained reflects recaptures of individuals with visible implants at each time interval accounting for the running total of individuals who lost implants. Actual percentage of retained implants is likely to be considerably higher than what is shown here. Two white-shouldered fairy-wren females were recaptured at the end of our study >27 months (823 and 828 days) after implantation, with implants ∼1/2 the original size.

## 4. Discussion

Exogenous testosterone studies have been integral to establishing which traits are under androgenic control. Though effects of pharmacological (i.e. unnaturally high) doses can reveal important information about hormone response mechanisms (Ketterson, 2014; Ketterson et al., 2005), physiologically-relevant manipulations are needed to establish which traits are sensitive to androgens under natural settings (Fusani, 2008; Goymann and Wingfield, 2014; Quispe et al., 2015). Delivering a physiological rather than pharmacological dose is an enduring challenge in hormone manipulation studies. We combined data collected across four separate implant studies using beeswax and peanut oil implants scaled to deliver 3 separate doses of testosterone. As reported in the initial paper introducing beeswax and peanut oil implants in female Japanese quail (Quispe et al., 2015), androgens peaked within a week of implanting and remained elevated for approximately 2 weeks (Figure 1). In captive zebra finch, 1 mg and 2 mg doses did not result in differences in plasma androgen levels. The dosage and time-release information we report here should provide a valuable reference for researchers aiming to deliver a physiological dose of testosterone to their respective study organisms, especially studies of small passerine bird species.

One proposed benefit of using implants composed of beeswax and peanut oil in wild organisms is that these materials should be readily dissolved, thus not requiring recapture and removal (Quispe et al., 2015). However, we found that implants were retained in many individuals for several months and even years in some cases, with two individual white-shouldered fairy-wrens captured with intact implants >2 years after implantation. At a bare minimum, ∼50% of implants were retained six months after manipulation, and >25% remained after one year (Figure 2). Documentation of long-term implant retention was not a motivation of any of the studies we combine here, so we only have information from opportunistic recaptures in each fairy-wren species. Hence the percentages we report in Figure 2 are surely an underestimation of how many individuals retained implants at each time point. In some cases of implants being retained for 6 months or more, implants were notably smaller than their initial size, but not fully dissolved (n = 5 of 16). That implants were diminished over time in these cases coupled with the fact that several individuals showed no implants within 6 months suggests that implants can be digested. Importantly, we excluded individuals who were suspected to have lost implants through improper sealing of the implant site, so individuals recaptured without implants presumably digested implants as expected. Given that wax is poorly water soluble, we suspect that the beeswax fraction of implants took the longest to degrade. Replacing beeswax with a more water soluble, but still moldable material should shorten implant degradation time.

We altered the fraction of beeswax to peanut oil slightly from 80:20 to 75:25 between field seasons in our white-shouldered fairy-wren study (Table 1). This was done to reduce crumbling of implants that often occurred when pinched with forceps during setting in our pilot season. Increasing the oil fraction of implants improved their solidity and we noted fewer cases of crumbling in our second field season. Because the delivered dose of testosterone is determined by the size and shape of the implant, any crumbling should result in implants being discarded and replaced to ensure delivery of the desired dose. We did not note any difference in dissolution of these two implant batches in the field, and due to inconsistent recapture and rare cases of dissolution we cannot adequately compare how different proportions of beeswax and peanut oil affected rate of dissolution (n = 1 dissolved implant of 7 recaptured from 80:20 beeswax:oil batch, and 4 dissolved of 23 recaptured from 75:25 batch).

## Conclusions

Beeswax/peanut oil implants are a cost-effective tool for delivering a scalable dose of hormones for around two weeks. However, our long-term field studies suggest the implants take considerably longer to fully degrade than previously suspected. While retained implants do not appear to affect physiology and behavior, their continued presence might pose a challenge for researchers interested in manipulating hormones in subsequent seasons. Replacing the beeswax fraction with a more readily digestible material should improve the rate at which implants degrade. Such modifications of the matrix should be validated for the hormone levels they result in. Implants should also be checked for hormone levels they produce in males and females as one cannot necessarily assume that sexes metabolize and excrete exogenous hormones in the same way. Continued innovation in hormone manipulation and delivery methods is important to optimize approaches that are cost-effective, minimally invasive, and mimic natural dynamics of hormone release as much as possible.

## Acknowledgements

We thank the many field and lab technicians across the four projects that comprise this study. For red-backed fairy-wren work, we thank field technicians Mary Margaret Ferraro, Maria Smith, David Weber, Sarah Duff and Malcom Moniz for their assistance in collecting samples in the field, and thank Southeast Queensland Water for access to our field site. The superb fairy-wren project was made possible thanks to field assistants Alessandro Turchi, Zachary Emery, Flavia Barzan, Anna Radkovic, Julia Kovacs, and Luke Ford. White-shouldered fairy-wren work wouldn’t have been possible without the support and hospitality provided by the Baivapupu clan in Western Province, Papua New Guinea. Funding came from the National Science Foundation (IOS-1354133 and IRES-1460048) and Tulane University Department of Ecology and Evolution (to S.K.) for red-backed fairy-wren work, and S.K. was supported by an NSF Graduate Research Fellowship during part of this work. Superb fairy-wren work was funded by the Holsworth Wildlife Research Endowment and the Ecological Society of Australia (to A.M/.), and the Australian Research Council (grant numbers FT10100505 and DP150103595 to A.P.). NSF IOS-1352885 and the American Ornithological Society (to J.B.) supported white-shouldered fairy-wren work. Finally, the zebra finch implant study was thanks to support from the Max Planck Society and R.Q. was supported by Convocatoria Nacional Subvencion a la Instalacion en la Academia 2021 N°SA77210088-ANID-Chile

## Supplementary materials

**Figure S1.**
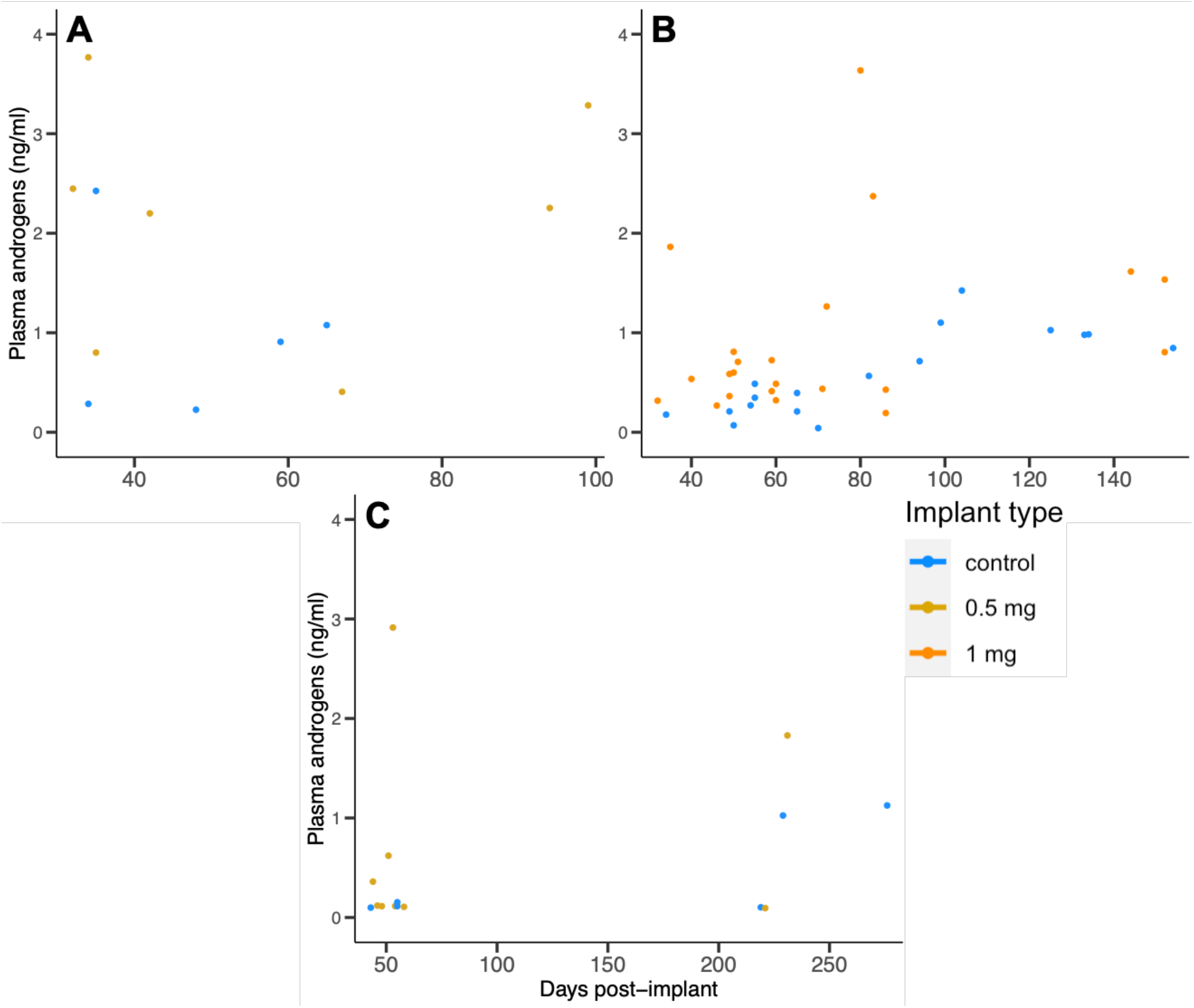
Time series of plasma androgens (ng/ml) measured after one month of implantation with a blank control or testosterone (approx. doses = 0.5mg and 1 mg) in **A**. red-backed fairy-wren (RBFW; n = 12 males), **B**. superb fairy-wren (SUFW; n = 39 males), **C**. white-shouldered fairy-wren (WSFW; n = 15 females). Individual hormone samples starting are shown across treatment groups absent loess lines due to discontinuous sampling.

